# Acquisition of hypoxia inducibility by oxygen sensing N-terminal cysteine oxidase in spermatophytes

**DOI:** 10.1101/2020.06.24.169417

**Authors:** Daan A. Weits, Lina Zhou, Beatrice Giuntoli, Laura Dalle Carbonare, Sergio Iacopino, Luca Piccinini, Vinay Shukla, Liem T. Bui, Giacomo Novi, Joost T. van Dongen, Francesco Licausi

## Abstract

N-terminal cysteine oxidases (NCOs) are enzymes that use molecular oxygen to oxidize the amino-terminal cysteine of specific proteins, thereby initiating the proteolytic N-degron pathway and thus conferring them oxygen-dependent instability. To expand the characterization of the plant family of NCOs (PCOs), we performed a phylogenetic analysis across different plant taxa in terms of sequence similarity and transcriptional regulation. Based on this survey, we propose a distinction of PCOs into two main groups: A-type and B-type sequences. A-type PCOs are conserved across all plant species and are generally unaffected at the mRNA level by oxygen availability. Instead, B-type PCOs differentiated in spermatophytes to acquire specific amino acid features and transcriptional regulation in response to hypoxia. Both groups of PCO proteins possess the ability to destabilize Cys-initiating proteins. Indeed, the inactivation of two A-type PCOs in *Arabidopsis thaliana*, PCO4 and PCO5, is sufficient to activate, at least partially, the anaerobic response in young seedlings, whereas the additional removal of B-type PCOs leads to a stronger induction of anaerobic genes and impairs plant growth and development. Our results show that both PCO types are required to regulate the anaerobic response in angiosperm. Therefore, while it is possible to distinguish two clades within the PCO family, separated by both amino acid features and transcriptional regulation, we conclude that they both contribute to restrain the anaerobic transcriptional program in normoxic conditions and together generate a molecular switch to toggle the hypoxic response in Arabidopsis.

**One sentence summary:** Hypoxic induction of Plant Cysteine Oxidases has been acquired and fixed in seed plants by ancestor proteins able to initiate the proteolysis of Cys-initiating protein substrates by the Arg/N-degron pathway.

## Introduction

The presence of an electron-rich sulfur sidechain makes cysteine residues extremely reactive and allows for a whole range of oxidative post-translational modifications (Reddie & Carroll, 2008). Some of these states are achieved through the initial oxidation of the free thiol group of cysteine, RSH, to RSOH (cysteine sulfenic acid). The RSOH state is very unstable and therefore can react with a second sulfenic group to generate a disulfide bridge, or RSOH oxidation can proceed to RSO_2_H (cysteine sulfinic acid). RSO_2_H may be further oxidized to yield the stable RSO_3_H state (cysteine sulfonic acid). Therefore, the cysteine oxidation reactions that yield cysteine sulfinic and sulfonic acid are usually considered as irreversible, while cysteine sulfenic acid may be reduced back to cysteine directly or via the formation of a disulfide bridge (Chung *et al*., 2013). Consequently, cysteine sulfenylation by reactive oxygen species (ROS) is responsible for changes in selective interaction, enzyme activity and substrate specificity of several proteins.

A specific case is represented by the oxidation of N-terminal cysteinyl residues to sulfinic acid, which has been shown to promote proteasomal degradation in plants and animal cells (Hu *et al*., 2005, Graciet *et al*., 2010). The sulfinic group supposedly mimics a carboxyl moiety of the glutamic and aspartic residues, which marks N-terminal degradation signals (N-degrons), following the pathway for proteolysis discovered by Varshavsky and colleagues (Bachmair *et al*., 1986). Here, N-terminally exposed oxidized cysteine provides recognition specificity by the N-terminal Arg-transferase (ATE) which catalyzes Arg conjugation. This N-terminal residue, in turn, is recognized by a single subunit E3 ligase Proteolysis 6 (PRT6) that marks the protein for proteasomal degradation with a chain of polyubiquitins (Graciet & Wellmer, 2010, Tasaki *et al*., 2012). Basal NO levels are also required to maintain the activity of this proteolytic pathway (Gibbs et al. 2014).

In plants and animals, N-terminal sulfinylation of specific proteins has been shown to be controlled by specific iron-dependent thiol dioxygenases, recently defined as N-terminal cysteine oxygenases (NCOs) (Weits *et al*., 2014, White *et al*., 2017, Masson et al. 2019). Since these enzymes use oxygen as a co-substrate, the N-degron dependent proteolysis of NCO substrates is promoted under oxic conditions and inhibited by hypoxia (Iacopino and Licausi, 2020, under revision). The first plant NCOs, Plant Cysteine Oxidases (PCOs) have been identified in Arabidopsis. Two of them, PCO1 and PCO2, have been initially identified among the proteins that constitute the core low-oxygen response (Weits *et al*., 2014). The requirement of iron for oxygen coordination in PCOs also generate a response to fluctuations in metal availability (Dalle Carbonare et al. 2019). Only few Arabidopsis proteins with N-terminal cysteine have been confirmed to be substrates of the N-degron pathway: the group VII Ethylene Response Factors (ERF-VIIs), the polycomb group protein Vernalization2 (VRN2) and Little Zipper Protein ZPR2 (Gibbs et al. 2011, Gibbs et al., 2018, Weits et al., 2019). However, oxidation of ZPR2 by PCO has not been tested yet. All these transcriptional regulators regulate adaptive responses to ambient and internal oxygen fluctuations (Giuntoli et al., 2017, Weits et al., 2020, Labandera et al., 2020). Recently, a PCO-counterpart of metazoans, the Cysteamine Dioxygenase ADO, was also found to regulate stability of Methionine-Cysteine initiating proteins (Masson et al. 2019).

The hypoxia-inducible PCOs are not the only member of this small family: for instance in Arabidopsis three additional proteins share sequence similarity with PCO1 and PCO2, which have been subjected to biochemical characterization *in vitro* (White *et al*., 2017, White et al. 2018). However, the role of the PCO that are not regulated by ERF-VIIs has not been elucidated *in vivo*, and the observation that they are not induced upon hypoxia suggests that they may function in different biological processes. In the present study, we filled this gap, by characterizing the role of the other PCO members with respect to hypoxia responses. We analyzed their evolutionary conservation by performing phylogenetic and functional analyses, and studied the impact of their genetic inactivation on plant physiology.

## Results

### Protein sequence and transcriptional regulation distinguish two conserved PCO clades

Whereas *PCO1* and *PCO2* are considered as part of the core anaerobic response genes in Arabidopsis (Mustroph *et al*., 2009), the transcriptional inducibility of the other three members has not been studied in detail. We therefore tested their mRNA level in response to low oxygen conditions (1% O_2_ V/V) in a time course over 12 h, corresponding to the light phase of the day. We confirmed that *PCO1* and *PCO2* are low-oxygen responsive, whereas no clear pattern of induction could be observed for the other three PCO genes (**Fig 1A**, **Supplemental Table S1**, **Supplemental Fig. 1-3**). However, previous analysis of microarray data showed that also *PCO4* is moderately induced during late hypoxia treatments (Weits *et al*., 2014). Therefore, to further verify whether hypoxic regulation is imposed at the transcriptional level, we generated a reporter line where the expression of the beta-glucoronidase (*GUS*) gene is fused to the upstream genomic region of the *PCO4* gene, including the intergenic sequence before the start of the transcription, the 5’ untranslated region (UTR) and the first 24 nt of the coding sequence (**Fig. 1b**). In seven-day old seedlings grown under aerobic conditions, GUS activity was observed in the vasculature, leaf primordia and basal zone of the first true leaves (**Fig 1b**). No increase in *PCO4prom:GUS* staining was observed after exposure to 6 h hypoxia in the dark, in either shoot or root tissues (**Fig 1b**). We therefore concluded that *PCO4*, similar to *PCO3* and *PCO5* is not induced by low oxygen conditions.

Next, we retrieved PCO-like sequences from angiosperm species for which the transcriptional response to low oxygen has been characterized at the whole-genome levels: Arabidopsis, rice (*Oryza sativa*), poplar (*Populus trichocarpa*), cotton (*Gossypium hirsutum*) and tomato (*Solanum lycopersicum*) (**Supplemental Table S2**). Since not all poplar sequences were represented on the microarray analysis available, we compared their expression level between aerobic and hypoxic (4 h 1% O_2_ v/v) conditions in a local poplar accession (*P. alba* ‘Villafranca’ clone). We aligned these putative orthologous amino-acid sequences and built a phylogenetic tree based on the conserved regions shared among them. The whole set of sequences separated clearly into two main clades: one (A-type PCOs) with proteins whose respective mRNA levels were not upregulated under low oxygen conditions, and a second clade (B-type PCOs) containing all low-oxygen inducible sequences (**Fig 1c**, **Supplemental File S1**). The two clades could be distinguished primarily due to the presence or identity of three different conserved regions within the ADO/PCO domain (interpro id IPR012864): a Glu/Asp acid triad at the beginning of the conserved (position 45-47 in AtPCO4), which is absent in the hypoxia-inducible PCOs, a substitution of Tyr with Phe/Leu towards the center of the ADO/PCO domain and the fixation of a Gly residue instead of an Ala/Thr within the highly conserved C-terminal part (**Fig. 1d**). This result hinted at a concomitant conservation of structural and cis-regulatory features for PCO genes in angiosperms.

**Figure 1.**
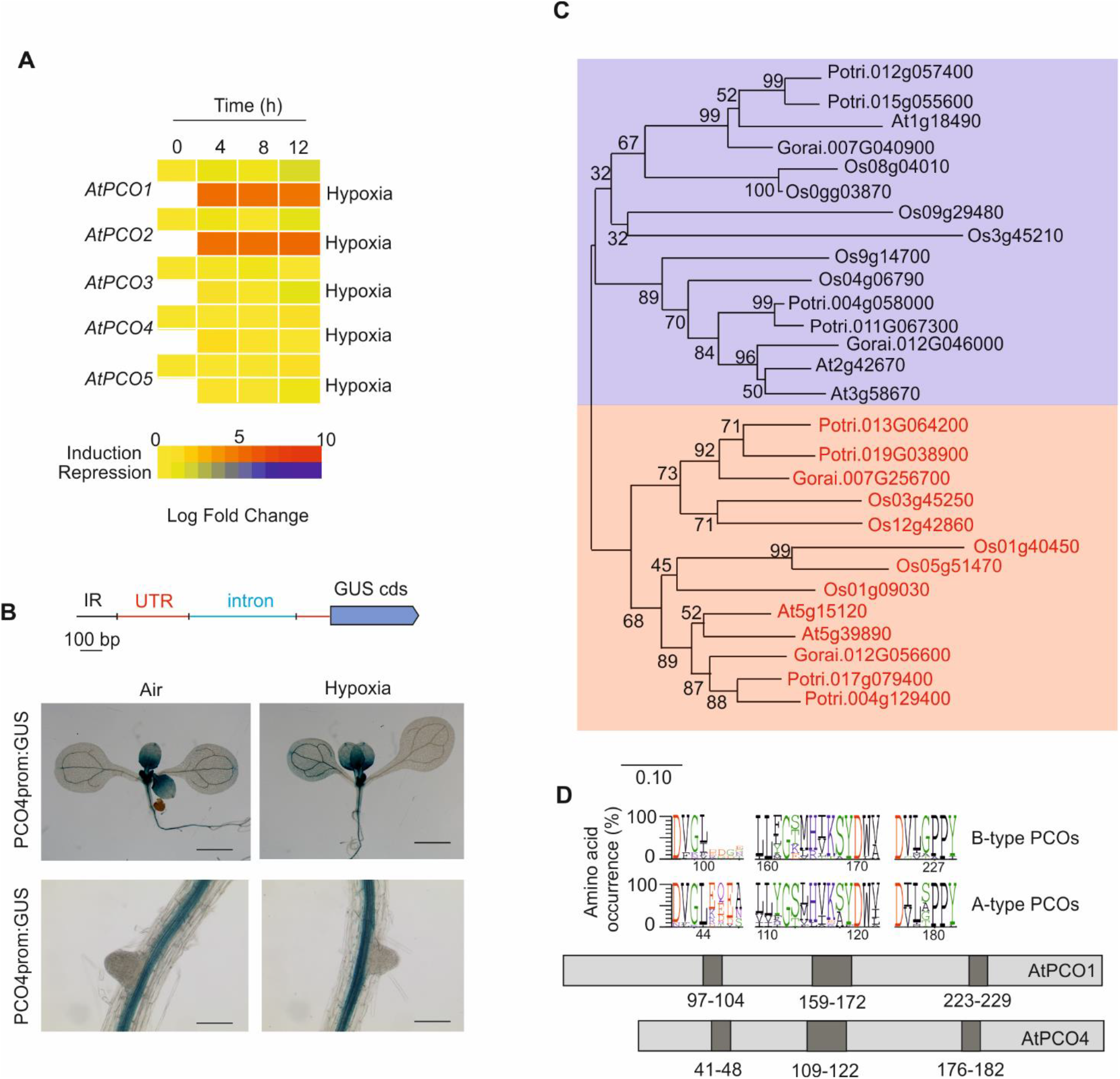
Hypoxic regulation of *PCO* genes in angiosperm species. **(A)** Expression analysis of the five PCO genes present in the Arabidopsis genome in response to hypoxia. mRNA was extracted from 4-day old seedlings treated for 0, 4, 8 and 12 hours of hypoxia (1% O_2_ v/v) and the expression of PCOs was compared to that of plants kept under normoxic conditions in the dark. Numeric expression values are shown in **Supplemental Table S1**. **(B)** Visualization of *PCO4* promoter activity by GUS-reporter staining in 7-day old seedlings treated with 6 h normoxia or hypoxia (1% O_2_ v/v). **(C)** Phylogenetic tree illustrating the relatedness of PCO proteins encoded by angiosperms for which transcript data about low oxygen responses are available: *Arabidopsis thaliana (At), Populus trichocarpa (Potri), Oryza sativa (Os), Solanum lycepersicum (Sl*), and *Gossipium raimondii (Gorai*). The Maximum likelihood method and JTT matrix-based model was used to generate an unrooted tree of PCOs. Branch length represent the number of substitutions per site. The percentage of trees in which the associated isoforms clustered together is shown next to the branches. Protein sequences used for the alignment are listed in **Supplemental Table S2**. The genes that encode for the PCOs that were shown to be significantly upregulated upon low oxygen exposure are indicated in red (p-value<0.05). **(D)** Amino acid occurrence in the three conserved regions that show the largest variation between A-type and B-type PCO clades. Their position in AtPCO1 and AtPCO4 protein is shown at the bottom of the panel. Letter height represent percentage of occurrence.

### The Hypoxia Responsive Promoter Element is conserved in the promoter of inducible PCO genes

The inducibility of B-type PCO genes by low oxygen conditions in the angiosperm species considered before is likely explained by the transcriptional regulation imposed by RAP2-type transcription factors, since this has been demonstrated in Arabidopsis previously (Gasch et al. 2016). Indeed, when analyzing the 1 Kb of the 5’ genomic sequence preceding the 5’ untranslated region (5’ UTR) of each hypoxia-inducible PCO-coding gene, we found one to four repeats of the Hypoxia Responsive Promoter Element (HRPE), identified as the main DNA feature recognized by the ERF-VII transcription factors (**Supplemental Table S2**). This 9 bp long motif occurred both in the 5’ untranslated region and the upstream intergenic region. Only in the case of the *Loc_Os1g09030* gene, the HRPE element was found inside a long 5’UTR, 1270 bp before the initial ATG codon (**Supplemental File S2**). We could not identify the same motif in the genomic region upstream of any of the A-type PCO genes. A chi-square analysis confirmed a significant correlation (P ≤0.001) between the occurrence of at least one HRPE element in the promoter or 5‘UTR and the regulation imposed by hypoxia. These results supported the hypothesis that genes coding for B-type PCOs are controlled by ERF-VII transcription factors.

### Hypoxia-inducible B-type PCOs have been acquired and conserved in spermatophytes

The identification of two separate clades of PCOs, namely A-type (not induced by hypoxia) and B-type (hypoxia inducible), led us to question when this speciation occurred during plant evolution, with possible implications in the mechanisms of oxygen perception in photosynthetic eukaryotes. We therefore searched for PCO-like sequences in genomes and transcriptomes of species belonging to taxa that could represent stepwise acquisition of traits that belong to actual angiosperms. To the angiosperm sequences used for the analysis shown in Fig. 1c before, we added those of *Pinus pinea* and *Picea abies* for gymnosperms, *Pteris vittata* and *Botrypus virginianus* for pteridophytes, *Selaginella moellendorfii* and *Lycopodium annotinum* for lycophytes, *Physcomitrella patens* and *Marchantia polymorpha* for briophytes. We also included three members of the green algae taxon: *Volvox carteri*, *Chlamydomonas reinhardtii* and *Dunaliella salina*. Reciprocal Blast-search using Arabidopsis PCOs as a reference showed that B-type sequences could only be found in spermatophytes, whereas evolutionary more primitive species only contained A-type proteins (**Supplemental Table S3**).

We expanded this initial analysis, by taking into consideration 212 PCO-like sequences from 42 plant species whose genome has been fully sequenced, including two belonging to the rodophyta and one from the glaucophyta taxa (**Fig. 2, Supplemental Table S3**). All sequences identified contain an ADO/PCO domain, characterized by highly conserved His residues in position 98, 100 (with respect to the PCO4 sequence) and, to a lesser extent, 164, which are involved in metal coordination and which are essential for dioxygenation catalysis (McCoy et al. 2006). Based on the three structural features identified above and depicted in **Fig 1d**, all PCO sequences from embryophytes could be distinguished in A-type and B-type. In few cases, highlighted in red in **Supplemental Table S3**, ambiguous attribution to either one of the two clades was solved by reciprocal blast against the Arabidopsis proteome. Algae PCO instead showed characteristics of both clades and were therefore considered as a separate one (**Fig. 2**, **Supplemental Table S3**). The highest number of PCO-coding genes was observed in monocots, fabales and *Malus domestica* among the rosales class, in agreement with the progression in genome size due to duplication events (Liu et al. 2016). Although the ratio B-type to A-type PCO varies considerably between species, at least one representative of both clades could be found in all spermatophytes considered. Among ferns and lycophytes, instead, only A-type PCOs were found, except for *Pteris vittata* which appeared to also contain a B-type PCO protein (**Fig. 2, Supplemental Table S3**).

**Figure 2.**
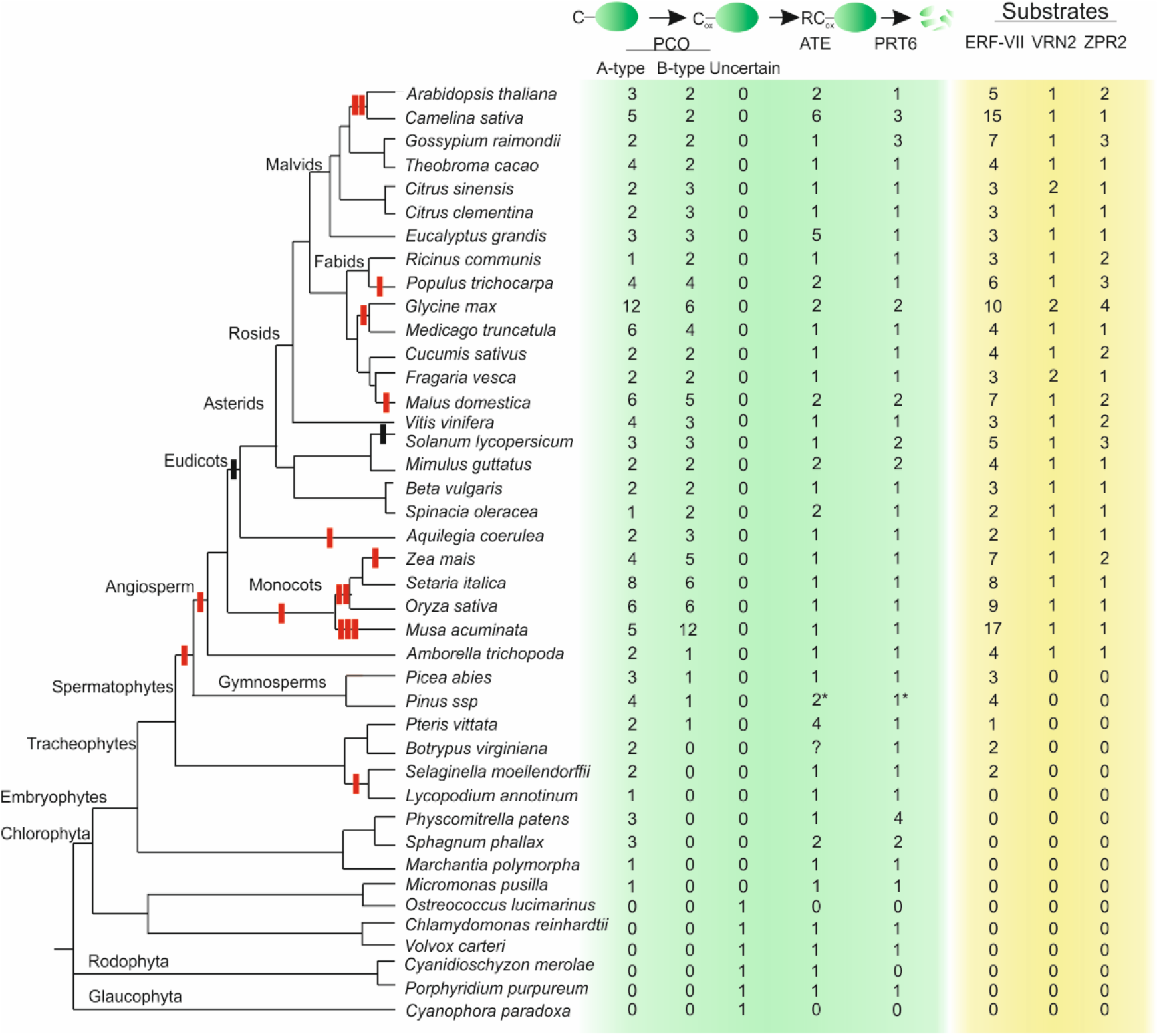
Evolutionary conservation across plant species of components of the oxygen-dependent branch of the N-degron pathway and their known targets. Whole-genome duplications are shown with vertical red bars for the branches corresponding to land plants, as described in **Liu et al (2016).** Vertical red ticks indicate genome duplication event, vertical black ticks represent genome triplication events. The number of non-identical sequences attributed to each class of predicted proteins and clade assignation for each PCO sequence is shown on the right side of the tree. The asterisks for ATE and PRT6 sequences in *Pinus spp* indicate that they were retrieved from the *Pinus tadea* predicted proteome, while PCO and ERF-VII sequences have been confirmed by cloning and sequencing from *Pinus pinea* mRNA. Question marks indicate identification of sequences that correspond to portions of the protein used as a bait but did not clear classification.

Finally, we interrogated public sequence databases to evaluate the co-occurrence of PCOs, the other two enzymes that act in the Arg/N-degron pathway, ATE and PRT6, and established Cys-degron substrates in the green lineage. We confirmed ubiquitous ATE and PRT6 presence in each examined species, although we could not always detect homologs in red algae, glaucophytes and green algae (**Fig. 2**). Cys-initiating proteins belonging to the ERF-VII group were identified in spermatophytes, including lycopods (**Fig. 2**, Supplemental Table S3), as previously indicated (Holdsworth and Gibbs, 2020). VRN2 and ZPR2 confirmed instead a later fixation in their Cys-initiating identity in angiosperms (**Fig. 2**, Supplemental Table S3Gibbs et al. 2019, Weits et al. 2020).

We therefore speculated that the A-type PCOs represent the earliest form of plant NCO, which acquired the ability to regulate the stability of a specific ERF group in vascular plants. Moreover, in spermatophytes the B-group PCO diverged from the original clade and acquired, at the gene level, ERF-VII-dependent inducibility through the HRPE motif.

### Chitryds are the only fungal species with PCO-like proteins

Since the existence N-terminal cysteinyl-dioxygenases (NCOs) has been confirmed in both plants and animals (Weits et al. 2014, Masson et al. 2019, Holdsworth and Gibbs 2020), we investigated their occurrence in the fungal kingdom. A thorough search throughout the proteome of fungi species whose genome has been fully sequenced returned hits only within the chytrid clade, although not all chytrid species tested showed a PCO-like sequence (**Fig. 3a, Supplemental Table S4, Supplemental Fig. 4**). Reciprocal identification of ADOs or PCOs in plant and metazoan databases using these fungal sequences as baits confirmed the orthology of the sequences. The presence of PCO/ADO-like sequences in almost all chytrid species in the database, but not in other phyla, can be explained as the loss of this enzymatic function in the fungal kingdom, while it was retained in chytrids. Alternatively, one can speculate about the acquisition of this enzyme exclusively by this latter fungal clade from a plant or metazoan host. We thus compared the most conserved regions identified in the ADO/PCO domain of species from the main phyla of the three kingdoms and used the resulting alignment to generate a phylogenetic tree with the Maximum Likelihood algorithm (**Fig. 3b**). Grouping of proteins reflected the assignation of original species into the three kingdoms, with chytrid putative NCOs clustering closer to plant PCOs than to metazoan ADOs (**Fig. 3b-c**). In light of this result, since speciation of Viridiplantae is estimated to have occurred before the separation of animals from fungi, we favoured the hypothesis of horizontal transfer of PCO-like gene from an ancestral green organism to a chitryd progenitor.

**Figure 3.**
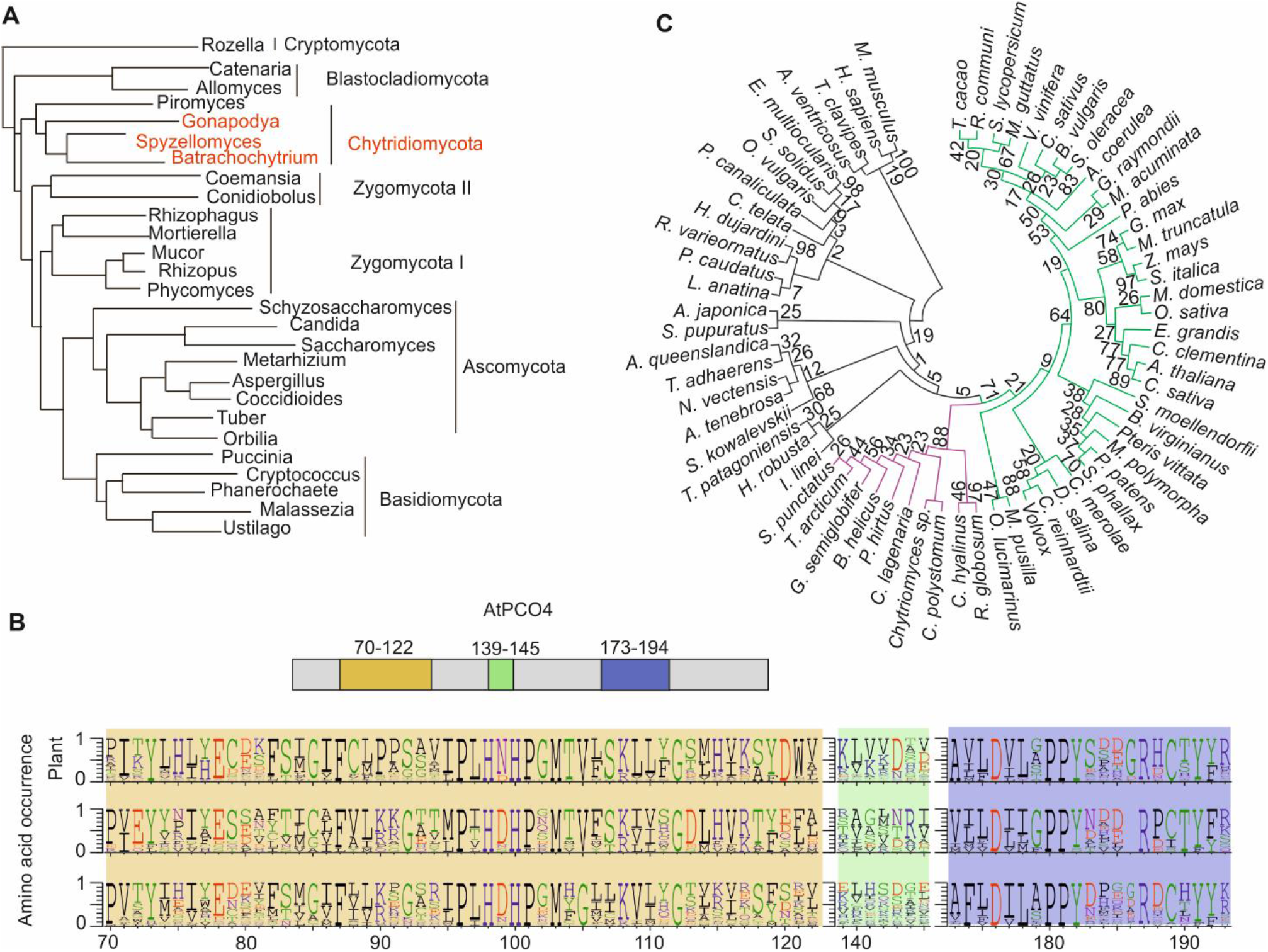
Presence of NCO-likes sequences across the fungal kingdom. **(A)** The phylogeny of fungi is based on the analysis by **Chang et al. 2015**. Species whose genome contains at least one NCO-like coding gene are marked in red. **(B)** Amino acid occurrence in three conserved regions of NCOs in animals, plants and fungi. Their position in AtPCO4 protein is shown at the top of the panel. **(C)** Relatedness of NCO proteins across the three eukaryotic kingdoms (plants in green, fungi in purple and animals in black), inferred by using the Maximum Likelihood method and JTT matrix-based model. Results of the bootstrap analysis are shown next to each branch.

### B-type PCO conservation is extended to the transcriptional regulation

Next, we investigated whether the genes encoding for B-type PCOs share conserved regulation by hypoxic conditions. Therefore, we quantified the mRNA levels of the PCO genes in one species of each taxon considered to generate the signature motifs depicted in **Fig. 1d**, selected depending on plant material available on site. Relative expression levels were monitored by realtime RT-qPCR, comparing mRNA extracted from samples exposed for different duration of hypoxic stress (1% O_2_ v/v). Control samples were maintained in the dark and harvested at the same time of the day, to disambiguate between possible circadian regulation and true hypoxic induction. High transcriptional up-regulation by hypoxia was observed for *PCO1* of *P. pinea*, which shares the strongest sequence similarity to the hypoxia-inducible B-type PCOs of angiosperm (**Fig. 4**, **Supplemental Table S5**). On the other hand, among more ancient species, moderate up-regulation was observed for *P. vittata PCO2* and *P. patens PCO2*, although mRNA of *P. patens* followed a similar trend under aerobic conditions in darkness (**Fig. 4**). PCO-like genes belonging to *S. moellendorffii* and *M. polymorpha* did not show altered expression in response to hypoxia. A search for HRPE-like elements in the 5’ upstream region of the *P. patens PCO2* gene did not lead to any positive identification, suggesting that its transcriptional regulation likely occurs via a different class of transcription factors or following an alternative mechanism (**Supplemental File S1**).

**Figure 4.**
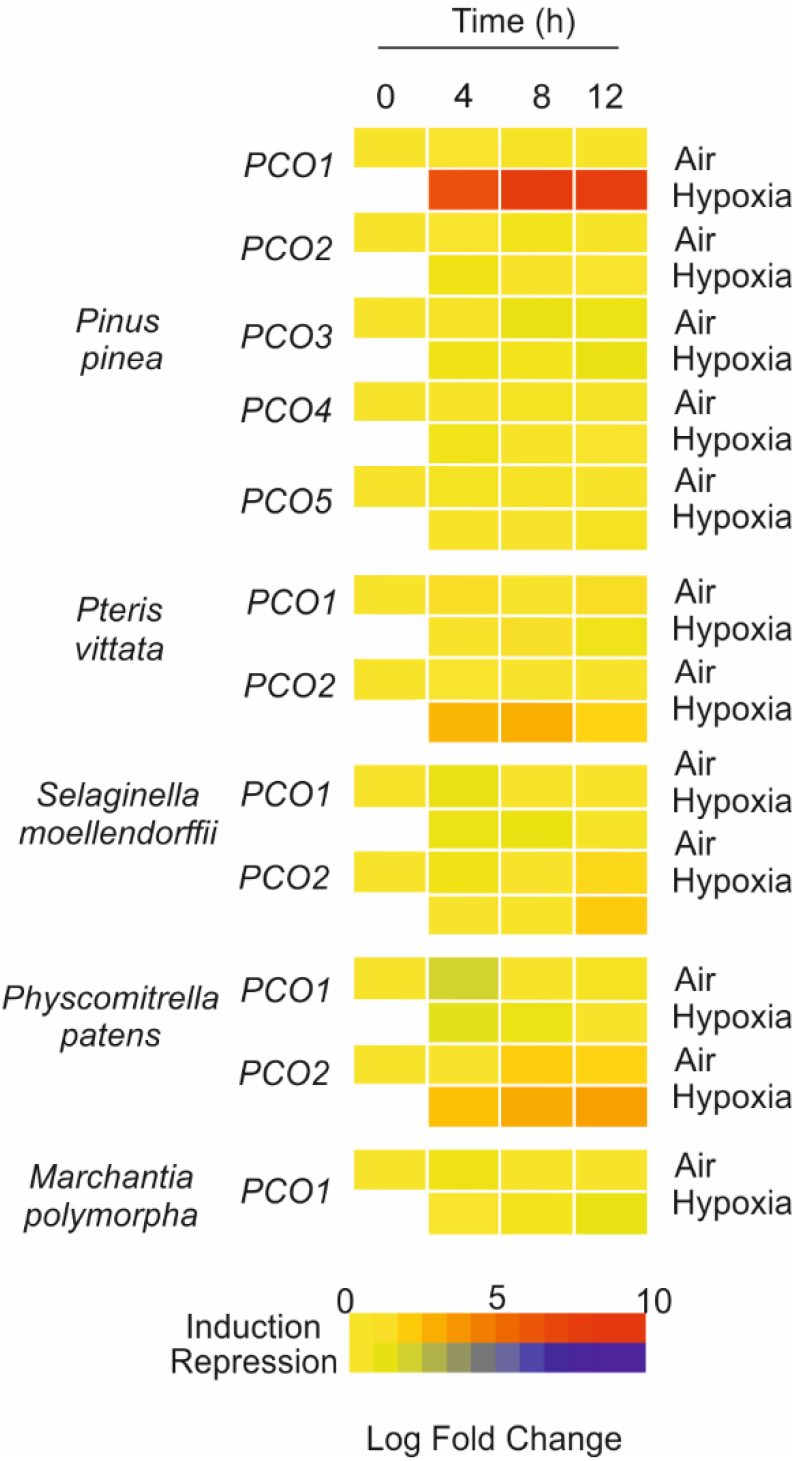
Expression analysis of PCOs in response to hypoxia in different plant species that represent subsequent steps in the evolution of land plants. mRNA was extracted from samples treated for 0, 4, 8 and 12 hours of dark hypoxia (1% O_2_ v/v) and the expression of PCOs was compared to that of plants kept under normoxic conditions in the dark (‘Air’). Numeric expression values are shown in **Supplemental Table S5**.

In conclusion, our results showed that early speciation of PCO from a common eukaryotic thiol dioxygenase ancestor occurred early in the plant lineage, while hypoxic induction by ERF-VII factors was only acquired in spermatophytes, and was accompanied by specific alterations in the amino acid sequence of the PCO proteins encoded by these genes.

### Non hypoxia-inducible PCOs contribute to control ERF-VII stability

Despite the structural differences highlighted in Fig. 3a, A-type and B-type PCOs share considerable sequence similarity among taxa, suggesting that their molecular function might be retained throughout evolution. B-type AtPCO1 and AtPCO2 have been shown to promote proteasomal degradation of ERF-VII proteins via the N-degron pathway *in vivo* and *in vitro* (Weits *et al*., 2014, White *et al*., 2017). We proceeded to analyze whether A-type AtPCO4 and AtPCO5 are also involved in the regulation of anaerobic responses, by controlling the stability, and thus activity, of proteins with a Cys-degron in an oxygen dependent manner. First, we tested the subcellular localization of PCO4 and PCO5: fusion to an N-terminal GFP showed nuclear and cytosolic localization, similar to what we have previously shown for PCO1 and PCO2 (**Fig. 5A**, (Weits *et al*., 2014)), independent of the occurrence of a region rich in positively charged residues typical of nuclear localization sequences (**Supplemental Table S6**). This result confirmed the potential of A-type PCOs to act on nuclear-localized transcriptional regulators. Moreover, heterologously expressed PCO4 and PCO5 protein consumed molecular oxygen when incubated in the presence of a five-amino acid long CGGAI peptide, which corresponds to the N-terminal consensus of ERF-VII transcription factors (**Fig 5B**), confirming their capacity to oxidize N-terminal cysteine, as observed *in vitro* (White et al., 2018).

**Figure 5.**
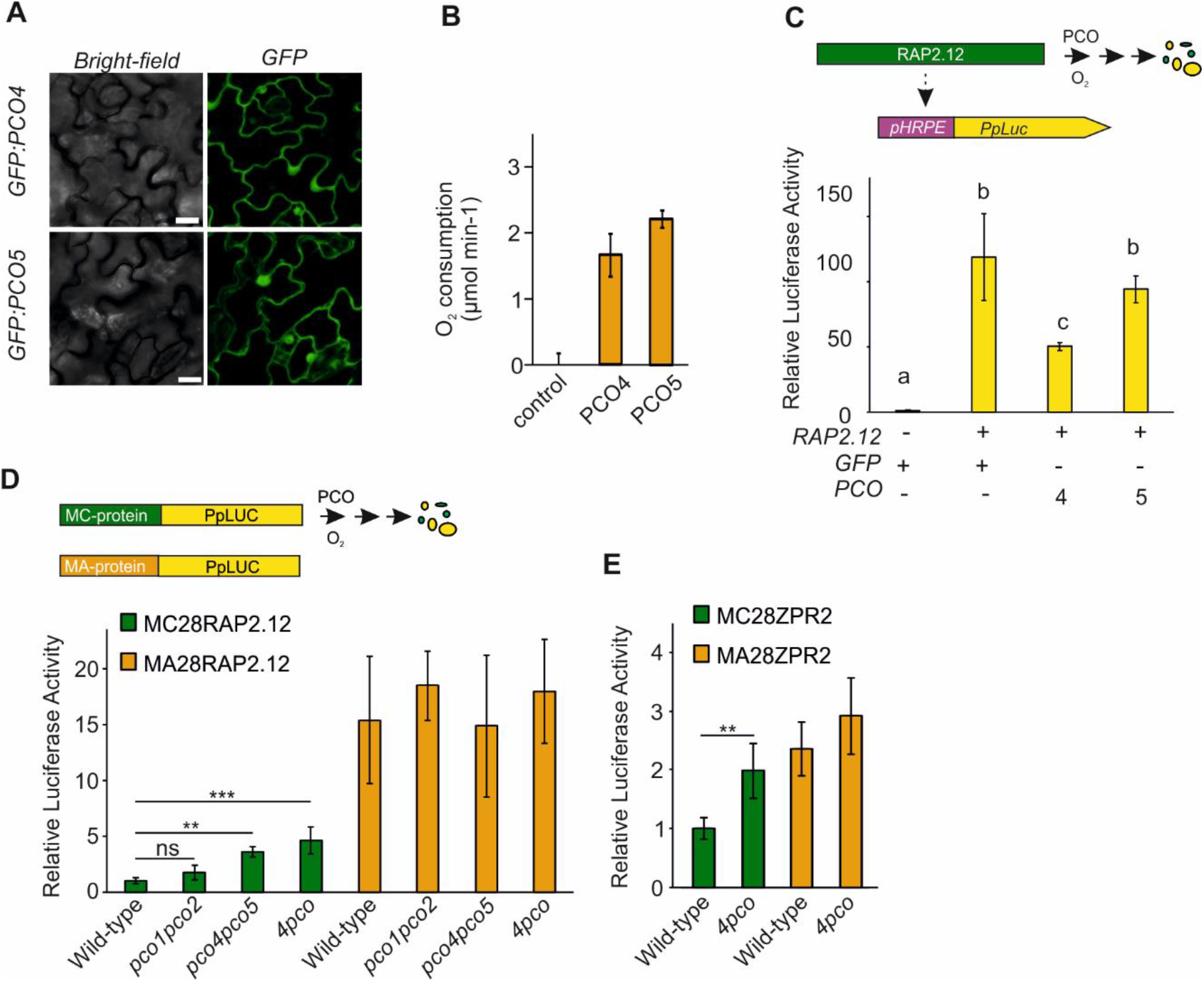
Role of A-type PCOs in controlling the activity of Cys-degron proteins. **(A)** Subcellular localization of PCO4 and PCO5 proteins harboring an N-terminally fused eGFP. Scale bars, 10 μm. **(B)** Oxygen consumption by purified PCO4 and PCO5 enzymes in a Cys oxidation *in vitro* assay using a CGGAI pentapeptide. Data are presented as mean±s.d. (*n*=6). Asterisks indicate statistically significant difference from a heat-inactivated protein extract as a control (*P*<0.05, one-way ANOVA). **(C)** Effect of *PCO4* and *PCO5* expression on transactivation imposed by RAP2.12 in *Arabidopsis thaliana* mesophyll protoplasts. To analyze RAP2.12 activity, a synthetic promoter harboring a five time repeat of the hypoxia responsive promoter element (HRPE) fused to a minimal 35S CaMV promoter driving expression of a firefly luciferase (*Photinus pyralis*) was used. A 35S:GFP vector was used as control. Data are presented as mean±s.d. (*n*=5). Letters indicate statistically significant difference (*P*<0.05, one-way ANOVA, Holm-Sidak post-hoc test). The experiment was repeated twice obtaining similar results. **(D** and **E)** N-degron reporter activity in plant genotypes with different PCO activity. Protoplasts of wild-type, double *pco1/2* and *pco4/5* mutants and quadruple *4pco* mutants were transiently transformed to express reporter (RAP2.12_1-28_-FLuc in **D** and ZPR2-FLuc in **E**) and normalizer luciferases (35S:RLuc). N-degron substrate reporters are shown in green, while stable reporters are shown in orange. Data are presented as mean±s.d. (n=5, the experiment was repeated twice). Asterisk indicate statistically significant difference (one-way ANOVA followed by Holm-Sidak post-hoc test in D, two-tailed t-test in E).

Next, we tested the ability of PCO4 and PCO5 to restrict ERF-VII activity *in vivo* by transient expression in mesophyll protoplasts. To this aim, we used the synthetic ERF-VII responsive promoter pHRPE (Kerpen et al. 2019). This was inserted in a reporter vector that bears two luciferase genes, a firefly (*Photinus pyralis*) luciferase under control of the HRPE promoter and a sea pansy (*Renilla reniformis*) luciferase driven by a 35S CaMV promoter, which was used for normalization purposes. We named this vector pHRPE-DL. Arabidopsis mesophyll protoplasts were co-transfected with pHRPE-DL, a construct designed to express the ERF-VII transcription factor RAP2.12 and either a vector bearing a PCO gene under control of the 35S promoter, or a GFP sequence as negative control. In this transient assay, PCO4 was able to repress RAP2.12 activity on the HRPE promoter, while PCO5 showed only a limited, statistically non-significant effect (**Fig 5C**).

Finally, we examined the effect of PCO on ERF-VII stability by using a chimeric reporter protein consisting of the N-terminal 28 aa of RAP2.12 fused to firefly luciferase. To this purpose, we selected Arabidopsis lines bearing a T-DNA within the transcribed sequence of *PCO4* and *PCO5*. We first identified homozygous *pco4* and *pco5* single mutants and subsequently crossed them to generate a double *pco4/5* knock-out mutant (**Supplemental Fig. S5**). For comparison, a previously identified *pco1/2* mutant was also included in the analysis. The activity of the chimeric reporter in protoplasts was enhanced in both *pco1/2* and *pco4/5* double mutants, and we observed an even stronger RAP2.12-PpLUC signal when this reporter was expressed in a *4pco* background where both A-type and B-type PCOs are knocked-out (**Fig. 5D**). This observation indicates that both PCO clades act redundantly to regulate ERF-VII proteolysis. We also observed enhanced stability of a full-length LITTLE ZIPPER 2 (ZPR2)-firefly luciferase fusion in the *4pco* mutant, providing evidence that ZPR2 is also a PCO substrate *in vivo*. The signal of MA-initiating versions of both chimeric reporters was higher than the MC version, and comparable between the wild-type and all *pco* knock-out genotypes (**Fig. 5E**). This confirmed that proteolysis initiated by PCOs requires an N-terminally exposed cysteine, while it also indicates that additional regulation occurs at the N-terminally exposed cysteine, possibly via the remaining PCO3 enzyme. Taken together, these observations support the hypothesis that A-type PCOs also possess the ability to oxidize N-terminal exposed cysteine and, in angiosperms, both clades act to restrict the anaerobic response by promoting proteolysis of substrates of the N-degron pathway.

### In Arabidopsis, A-type PCOs play a role to repress the hypoxic response under aerobic conditions

Previously, we showed that PCO1 and PCO2 repress the hypoxic response under aerobic conditions (Weits et al., 2014). Since also PCO4 and AtPCO5 are able to promote ERF-VII degradation, their inactivation would be expected to lead to an induction of the hypoxic response. We therefore aimed at discerning the contribution of each PCO clade on the regulation of the anaerobic response, by analyzing the expression of seven genes that belong to the core anaerobic response (Mustroph et al. 2009) in the *pco1, pco2, pco4, pco5, pco1/2,pco4/5* and *4pco* mutants and compared their expression to that of wild type plants. The selected genes included those involved in fermentation (*Alcohol Dehydrogenase ADH, Pyruvate Decarboxylase PDC1, Sucrose Synthase SUS1* and *SUS4* (Santaniello *et al*., 2014)) and signaling (*Hypoxia Responsive Attenuator 1 HRA1* (Giuntoli *et al*., 2014) and *LOB Domain transcription factor 41 LBD41*). In adult 4-week old plants we could observe only a minor and statistically non-significant effect on the expression of these genes in the *pco1/2* and *pco4/5* mutants. Instead, the *4pco* mutant showed a significant increase in the expression of the anaerobic genes when compared to the wild-type, showing that both PCO clades act redundantly under aerobic conditions at this developmental stage (**Fig. 6A**). Comparable anaerobic gene expression was observed in *pco* mutant lines using 10-day old plants, whereas 5-day old *pco1/2* and *pco4/5* seedlings showed increased hypoxia-inducible transcripts under aerobic conditions, albeit not as strong as in the *4pco* background (**Fig. 6A**). These observations show that both PCO groups are involved in the repression of anaerobic responses and confirm the existence of age-dependent regulation imposed on the activation of anaerobic genes, as reported before (Giuntoli et al. 2017).

**Figure 6.**
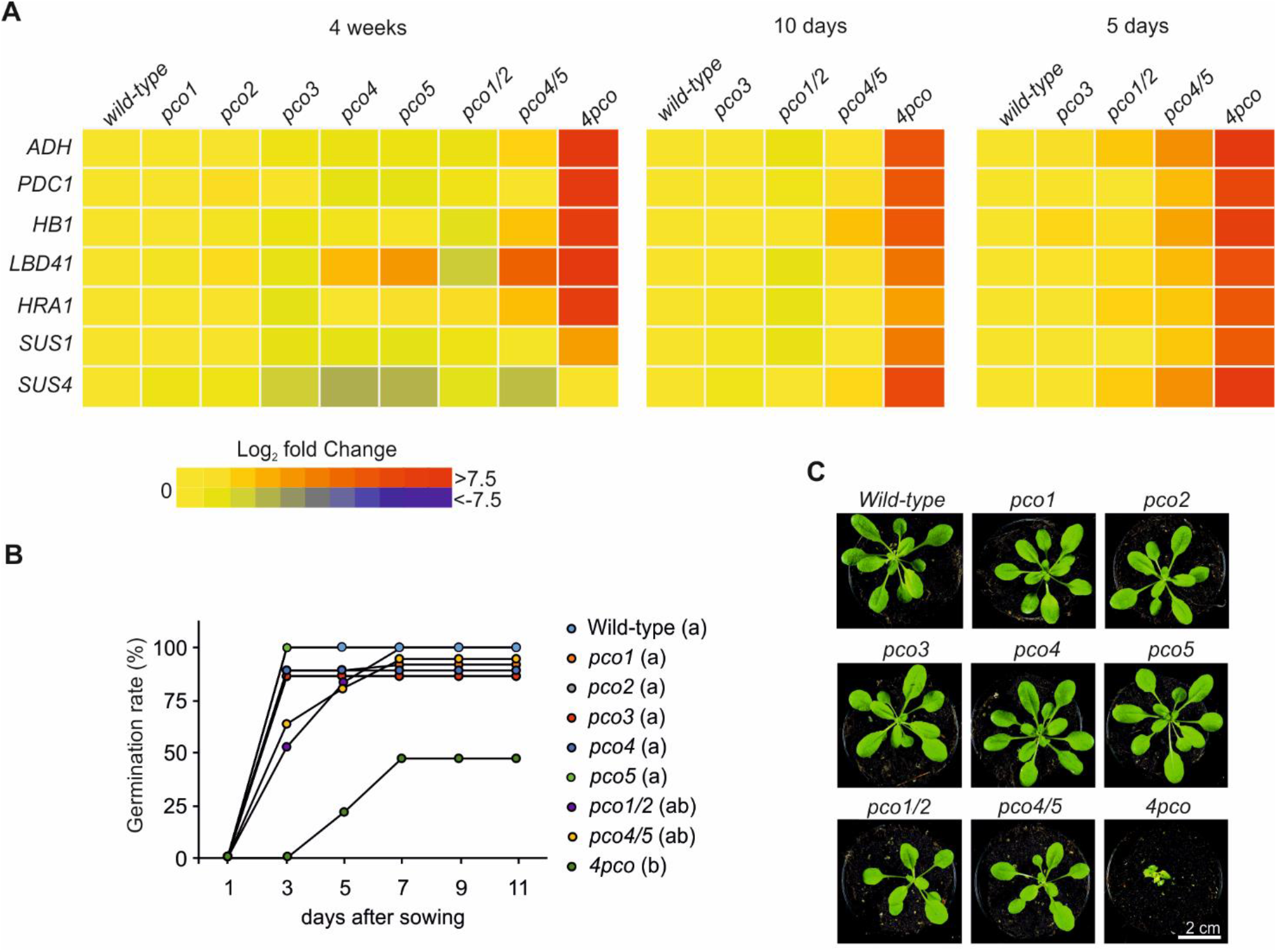
The effect of PCO knock out mutants on plant growth and the molecular response to anaerobiosis. (A) Phenotype of pco4, pco5 and pco4pco5 mutants grown on agarized MS medium. Scale-bar (A) Heat map depiction of the transcript levels of 10 genes belonging to the core set of the anaerobic response in single (*pco1, pco2, pco3, pco4* and *pco5*) double (*pco1/2* and *pco4/5*) and quadruple (*4pco*) mutants grown for four weeks, 10 days and 4 days under normoxic conditions. Numeric expression values are shown in **Supplemental Table S7.** (**B**) Germination percentage of wild-type and *pco* mutant seeds. Letters close to the genotype legend indicate grouping according to statistically significant differences. This was assessed by Kaplan-Meier Survival Analysis (Log-Rank) followed by Holm-Sidak post-hoc test. (C) Phenotype of wild-type and *pco* mutants grown in soil for four weeks. Scale bar, 1 cm.

Since expression of hyper-stable versions of ERF-VII and ZPR2 caused altered plant phenotype (Giuntoli et al. 2017, Weits et al. 2020), we also characterized the consequences of PCO inactivation on development and growth. When grown in vertical plates, the high-order *pco* mutants exhibited delayed germination although this reduction was significant only for the *4pco* genotype, while the single mutants could not be distinguished from the wild type (**Fig. 6B**). Moreover, when grown in pots, *4pco* mutant adult plants developed wrinkled, pale leaves characterized by a higher degree of serration (**Fig. 6C**), while the single and double mutants remained undistinguishable from the wild type. Altogether, the gene expression and phenotypic analyses support a high degree of redundancy among PCO isoforms in controlling the anaerobic response in plants and preventing the developmental consequences of its activation under aerobic conditions.

### Constitutive induction of anaerobic genes in *pco4pco5* mutants affects anaerobic survival

Constitutive activation of the anaerobic response was shown to have a negative effect on the overall ability of plants to endure actual oxygen deficiency, depending on the experimental conditions employed (Licausi *et al*., 2011, Gibbs *et al*., 2011, Riber *et al*., 2015). Keeping this in mind, we tested the relevance of PCO4 and PCO5 in tolerance to temporary oxygen deficiency. We did not include the quadruple *4pco* mutant in this survey, since its pleiotropic phenotype made it extremely difficult to obtain a sufficient number of homogenous plants to be treated, and its altered growth hindered proper fitness scoring (Masson et al. 2019, **Fig. 6C**). Thus, we first compared the anoxic survival rate of single *pco4* and *pco5* and double *pco4/5* mutants with that of the wild type when plants were grown on a sugar-supplemented medium. In this condition, the double mutant exhibited a significant improvement in anoxic tolerance (**Fig. 7A and B**), possibly due to a primed state of acclimation to anaerobic conditions in the presence of sufficient supply of carbon for glycolysis and fermentation. Indeed, in a separate test conducted with exogenously supplemented sucrose, the *pco4/5* mutant exhibited high fermentative potential under aerobic and hypoxic conditions. In particular, it produced in air as much ethanol as the wild type after 12 h of hypoxia (**Fig. 7C**). On the other hand, we observed the opposite trend when we exposed soil-grown plants to flooding as a mean to impose oxygen deprivation: the double mutant was significantly reduced in terms of biomass production after a four-day submergence in the darkness (**Fig 7D-E**). We interpreted this result as a negative effect of enhanced fermentation when plants experience hypoxia in conditions of severe carbon limitation.

**Figure 7.**
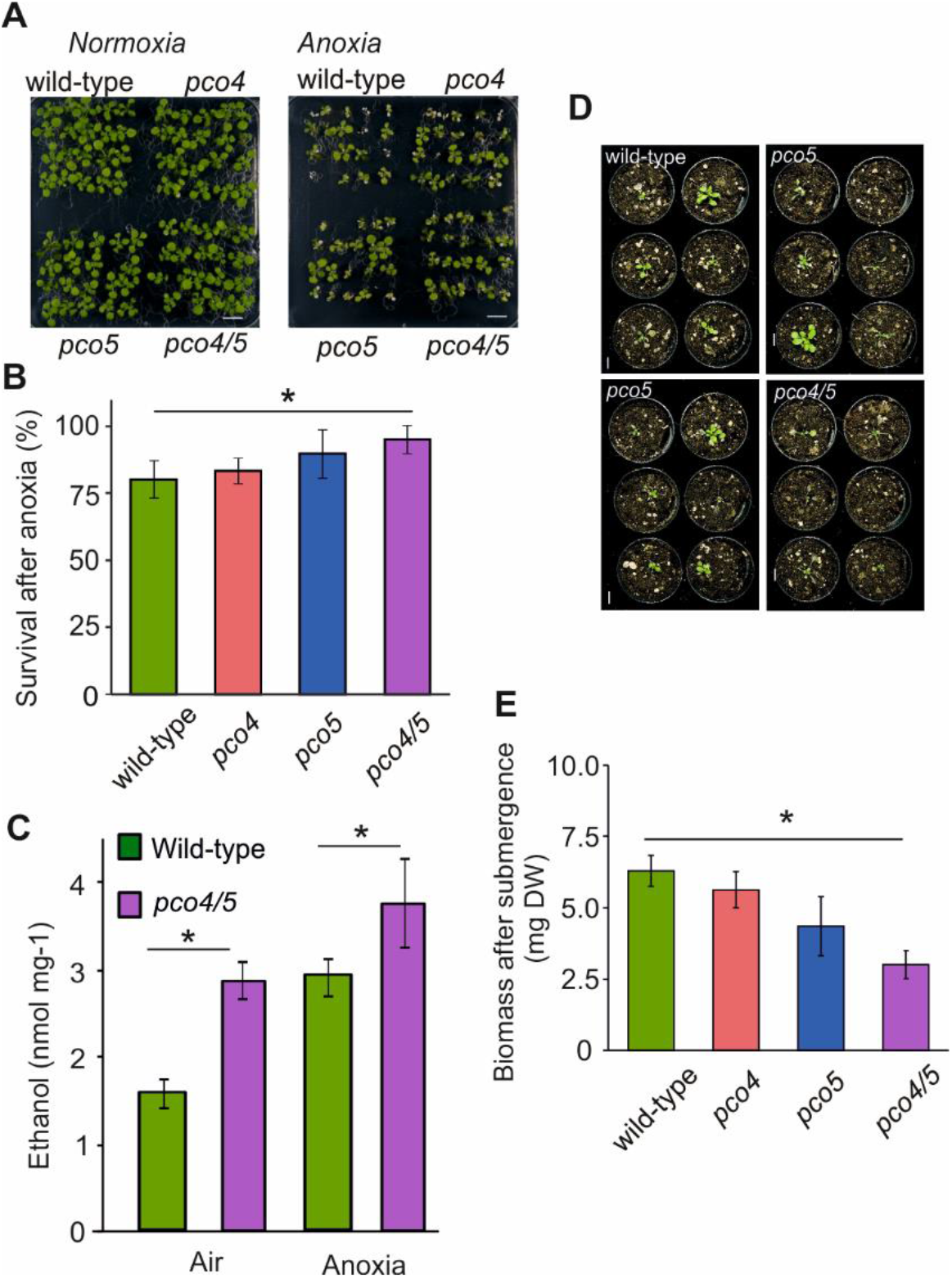
Contribution of PCO4 and PCO5 to the tolerance of Arabidopsis plants to low oxygen conditions. (**A**) Phenotypes of in vitro-grown wild-type, *pco4*, *pco5* and *pco4/5* plants 7 d after exposure to 9 h of normoxia or anoxia (<0.01% O_2_ v/v). Scale bar 1 cm. (**B**) Percentage of wild-type, *pco4, pco5* and *pco4pco5* plants surviving 9 h of anoxia in the dark, following a 7 d recovery period. (**C**) Ethanol production in tissues of wild-type and the double *pco4/5* mutant. One-week old plants were transferred to sterile media containing 2% (w/v) sucrose 12 h before being incubated for 12 h with anoxia or normoxia. The box plots are built on the basis of five replicates. (**D**) Phenotypes of soil-grown wild-type, *pco4, pco5* and *pco4pco5* plants recovering from 96 h of complete submergence in the dark. Scale bar 1 cm. (**E**) Biomass (dry weight) of wild-type, *pco4, pco5* and *pco4/5* plants following 7d of recovery from 96 h of dark submergence. Asterisks indicate statistically significant difference from wild type (P<0.05, one-way ANOVA).

## Discussion

In the present study, a phylogenetic analysis of the occurrence of NCO sequences among eukaryotes revealed that these are ubiquitously present in the plant kingdom (**Fig. 2**), as they are in the animal one (Masson et al. 2019). Sequences with high similarity to plant cysteine oxidases were fixed in embryophytes, possibly already to catalyze N-terminal cysteine oxidation, where this family proliferated due to genome duplication events. Among the classes of Cys-starting proteins identified so far in plants as PCO substrates, ERF-VIIs seem to be the earliest to be fixed in vascular plants, while ZPR2 and VRN2 proteins followed, in angiosperms (**Fig. 2**). This suggests a progressive co-adaptation with PCO to accommodate the N-terminal degron to PCO activity. It is tempting to speculate about the requirement for an ERF-VII/PCO circuit when plants acquired with vascularization a level of complexity that entails internal oxygen gradients (van Dongen and Licausi, 2015). Hypoxia-responsiveness is not a conserved feature of these proteins, which we defined here as A-type PCOs. Our analysis indicates that, instead, hypoxia-inducible B-type PCOs represent a relatively recent acquisition by spermatophytes, where a second group branched out from the original family (**Fig. 1C**). The innovation of these B-type PCOs is not limited to the protein sequence, where we could identify three main signature motifs that distinguish A- and B-type PCOs (**Fig. 1D**), but also extends to the acquisition of a DNA element, in the promoter of their respective genes, that confers ERF-VII mediated regulation under hypoxia (**Supplemental Table 3**). The conserved co-occurrence of this cis-regulatory feature and structural characteristics is suggestive of an optimization of anaerobiosis-inducible isoforms to reduced oxygen availability. In this way, B-type PCOs may play a role during hypoxia to specifically recognize ERF-VII proteins to ensure efficient and rapid restraint of anaerobic responses such as fermentation, whose excess has been shown to be detrimental for plant survival under submergence (Licausi et al. 2011, Paul et al. 2016). Remarkably, the need of a similar feedback loop has also been reported for mammals. Here, the oxygen sensing enzyme Prolyl Dehydrogenases (PHDs) control the stability of the α subunit of the hypoxia-inducible factor-1 (HIF1) complex, which, in turn, further upregulates PHD expression (Henze and Acker, 2010).

As mentioned above, NCO seem to be ubiquitously distributed and conserved in the plant and animal kingdom. Fungi, on the other hand, represent a peculiar case: we could only find NCO-like sequences in chytridiomycota, one of the early diverging lineages of this kingdom, but not in other groups (**Fig. 3A**). Not all chytrid species tested possess a PCO-like sequence, suggesting that this gene is not essential for the biology of these organisms (**Supplemental Fig. S4**). The fact that chytrids thrive in aquatic habitats, and water is necessary for the movement of chytrid zoospores, support the hypothesis of NCO-like sequences being involved in oxygen sensing in these organism (Kagami et al. 2014). In this perspective, NCO-mediated responses might help to cope with oxygen fluctuations in acquatic environments. Future characterization of fungal NCOs and their substrates, both *in vivo* and *in vitro*, will shed light on these aspects.

The absence of NCO-like sequences in all other groups might indicate that this has been lost early during fungal evolution, except than in chytrids, possibly because it affected fitness negatively. However, expression of plant PCO or human ADO in the budding yeast *Saccharomyces cerevisiae* did not impair cell growth, opposing the idea of toxicity of this gene in fungi (Puerta et al. 2019, Masson et al. 2019). Alternatively, horizontal gene transfer might have led to NCO fixation in chytrid genomes. Indeed, several chytrids parasitizes animal or plants species. The closest similarity of chytrid NCOs to PCOs rather than animal ADOs would favor the hypothesis of an acquisition from a plant donor (**Fig. 3B-C**). In support of this, chytrid fossils from the Rhynie chert, a Devonian-age lagerstätte, are parasites of rhyniophytes, demonstrating early establishment of plant parasitism (Taylor et al. 1994).

As mentioned above, the A-type PCOs do not exhibit conserved responsiveness to hypoxia at the transcript level. Nevertheless, PCO4 has been confirmed as a potential initiator of the proteolytic N-end rule pathway *in vitro* (White *et al*., 2017) and indeed could repress RAP2.12 activity in a transient assay, and inhibit the accumulation of N-degron reporters (**Fig. 5C-E**). The observation that a concomitant knocking-out of *PCO4* and *PCO5* genes induces activation of the anaerobic response in young Arabidopsis seedlings (**Fig. 6A**), indicates that their affinity for ERF-VII is sufficient *in vivo* to control the stability of these TFs under aerobic conditions. On the other hand, at later stages, the remaining PCO enzyme in Arabidopsis, or parallel repressive mechanisms, were sufficient to repress ERF-VII activity and thus induction of anaerobic genes (**Fig. 6A**). We reported similar observations for other mutants of the N-degron pathway, whose ability to further process substrates with N-terminal Cys is impaired (Giuntoli et al. 2017). Remarkably, while loss of four out of five PCO enzymes deregulates the anaerobic response, all other genotypes analyzed so far displayed some degree of complementation, suggesting the establishment of additional mechanism that restrict ERF-VII activity some days after germination. It is also interesting to note that all four PCO sequences analyzed so far can enter the nucleus (**Fig. 5A**, Weits *et al*., 2014) irrespective of the presence of typical nuclear localization sequences, suggesting that PCOs may be imported into the nucleus by a different mechanism.

Assuming that one of the roles of the PCOs is to restrain the activation of fermentation under aerobic conditions, it is not surprising that their absence negatively affects plant fitness, as seen by a decrease in germination speed and shoot growth in the *4pco* mutant (**Fig. 6B-C**). It is likely that these plants are exposed to a similar metabolic stress as that identified in plants expressing a stabilized version of RAP2.12 (Paul *et al*., 2016). Additional explanation for the *4pco* phenotype may be found via PCOs regulation on cys-degron proteins VRN2 and ZPR2, which are *in vitro* and *in vivo* PCO substrates respectively (Gibbs et al., 2018, **Fig. 5E**). VRN2 and ZPR2 regulate different aspects of plant development and accumulate in meristematic niches, tissues that have been characterized as chronically hypoxic (Gibbs et al., 2018, Shukla et al., 2019, Weits et al., 2020). Excessive abundance of these MC proteins in *4pco* mutants may therefore also play a role in causing its peculiar phenotype.

When single or double *pco* mutants, which do now show remarkable morphologic alterations under non-stress conditions, are challenged with oxygen deprivation, their performance strongly depends on the availability of carbon resources to support energy production via glycolysis coupled to fermentation. Indeed, double *pco4/5* mutants showed increased survival rate to anoxic exposure when grown, in the presence of exogenous sucrose (**Fig. 7A-B**), whereas they exhibited impaired biomass maintenance when grown in soil and subjected to prolonged submergence (**Fig. 7D-E**). In the past, contrasting results have been obtained when comparing plants impaired in ERF-VII degradation and subjected to oxygen deprivation (Gibbs *et al*., 2011, Licausi *et al*., 2011). Most divergent outcomes have been reported in the case of submergence, which is a compound stress that involves several factors in addition to reduced oxygen availability (Bailey-Serres & Colmer, 2014). An extensive analysis of the possible reasons for this have been carried out by Riber and colleagues, pointing at the importance of humidity in post-submergence conditions and the content and usage of carbon reserves in the plant (Riber *et al*., 2015). Due to the high relevance of this topic for crop breeding and farming practices, most useful and conclusive information on this matter is likely to come from the analysis of plant performance, when grown and challenged with submergence in proper agricultural conditions i.e. in open fields.

## Conclusions

The study presented here allows for a subdivision of PCOs into two clades based on their amino acid sequence and their transcriptional regulation: those that are induced upon hypoxia (B-type) and those whose expression is unaffected by oxygen limitation (A-type). Both PCO clades are involved in the repression of the anaerobic response under normoxic conditions. Interestingly, A-type PCOs are present ubiquitously, whereas the B-type enzymes have evolved with spermatophytes. In gymnosperms and angiosperms, the hypoxia-inducible clade PCOs may therefore have evolved as a mechanism to fine-tune the extent of the anaerobic response to the strength of the hypoxic stress. The Arabidopsis proteome contains over 200 proteins with a cysteine in amino terminal position. These proteins are all potentially targets of PCOs and their identification may lead to more information on which other processes may be regulated by the oxygen-dependent branch of the N-degron pathway for protein degradation.

## Materials and methods

### Plant material and growth conditions

*Arabidopsis thaliana* Columbia-0 (Col-0) was used as wild-type ecotype. Single *pco4* (GABI_740F11) knockout seeds were obtained from GABI-Kat. Single *pco5* (SALK_128432) knockout seeds were obtained from the Nottingham Arabidopsis Stock Centre (NASC). Homozygous lines were identified via PCR screening of genomic DNA using gene-specific primers together with T-DNA-specific primers. Double homozygous lines were obtained by crossing the two single mutants and then screening the F2 generation as described above. *Pinus pinea* nuts were collected along the Arno river in a 300 m^2^ area centered around Google maps coordinates 43.704733, 10.424682. *Pteris vittata* spores were collected from spontaneous plants found in the garden surrounding Casa Pacini of the Department of Crop Plants of the University of Pisa (coordinates 43.704733, 10.424682). *Selaginella moellendorfii* was purchased by Bowden Hostas and propagated vegetatively. *Physcomitrella patens* was provided by Tomas Morosinotto (University of Padova). *Marchantia polymorpha* Cam2 was provided by Linda Silvestri (University of Cambridge).

Growth in soil of Arabidopsis plants: seeds were sown in moisted soil containing pit and perlite in a 3:1 ratio, stratified at 4 °C in the dark for 48 h and then germinated at 22 °C day/18 °C night with a photoperiod of 8 h light and 16 h darkness with 80-120 μmol photons m^−2^s^−1^ intensity. For qPCR experiments on adult plants, 5-week-old plants were treated with low-oxygen in plexiglas box flushed with an artificial atmosphere containing nitrogen and oxygen in the proportions defined in the results section.

Axenic growth of Arabidopsis plants: seeds were sterilized using 70% ethanol for 1 min, incubated in 0.04% bleach for 10 min and rinsed six times with 1 ml distilled sterile water. Seeds were resuspended in 1 ml sterile water. Growth in liquid medium was performed inoculating 100 μl of seed suspension corresponding to 20-40 seeds in 2 ml of sterile MS medium (basal salt mixture, 2.15 g l-1, pH 5.7) supplemented with 1% sucrose in each well of 6-well plates. Seeds were incubated in the dark at 4°C for 48h and subsequently germinated for four days at 22 °C day/18 °C night with a photoperiod of 12 h light and 12 h darkness. Growth on solid medium was performed in square dishes (10 cm side) containing 40 ml of solid MS medium (basal salt mixture, 2.15 g l^−1^, pH 5.7) supplemented with 1% sucrose and 0.8% Agar. After stratification for 48h at 4°C in the dark, germination and growth of the plants occurred at 22 °C day/18 °C night with a photoperiod of 12 h light and 12 h darkness.

*Pinus pinea, Pteris vittata* and *Selaginella moellendorffii* were grown on perlite soil under growth chamber conditions as described above for *Arabidopsis thaliana*. Plantlets of *Populus alba* ‘Villafranca’ clone were maintained in in vitro conditions and treated with hypoxia as described in Dalle Carbonare et al. 2019. *Physcomitrella patens* was cultured in sterile conditions on solid Knop medium described in (Reski & Abel, 1985) while *Marchantia polymorpha* in solid MS half-strength medium (0.9% w/v Agar).

### Low oxygen treatments

Plants were subjected to low oxygen treatments inside Plexiglas boxes where an artificial atmosphere containing a mixture of oxygen and nitrogen gases according to the ratios defined in the results session was continuously flushed. During the hypoxic treatments, the boxes were maintained in the dark to avoid oxygen release by photosynthesis. Individual boxes were used for each time point to avoid cycles of hypoxia and reoxygenation. Plants used for control samples were maintained in the dark for an equal amount of time. Survival rate based on emergence of new leaves was measured seven days after the exposure to anoxic conditions. Submergence was imposed to four-week old plants grown in soil as described above, inside glass tanks entirely covered with aluminum foil to maintain dark conditions. Deionized water was equilibrated for 12 h to the room temperature before pouring it slowly into the tanks up to a 15 cm from the bottom of the tank.

### Cloning of the plant and bacterial expression vectors

Coding sequences and promoters were amplified from complementary cDNA or genomic DNA templates respectively using the Phusion High Fidelity DNA-polymerase (New England Biolabs). RAP2.12_1-28_-LUC and ZPR2-LUC gene fusions were obtained by overlapping PCR. All open reading frames were cloned into pENTR/D-TOPO (Thermo-Fisher Scientific). The resulting entry vectors were recombined into destination vectors using the LR reaction mix II (Thermo-Fisher Scientific) to obtain the novel expression vectors. A complete list and description of the oligonucleotides and destination vectors used is provided in **Supplemental Table S7** and **S8**, respectively.

### Identification of NCOs

Identification of NCO protein sequences in different sequenced plant species was performed by searching the phytozome database (www.phytozome.net). *Pteris vittata* and *Pinus pinea* sequences were retrieved from the 1000 Plants transcriptome database (www.onekp.com) and EuropineDB (http://www.scbi.uma.es/pindb/), respectively. Protein sequences similar to *Arabidopsis thaliana* PCO1 were retrieved using the BLAST algorithm (Atschul *et al*., 1990). The sequences obtained in this way were subsequently aligned back against the *Arabidopsis thaliana* protein database to ensure that they represent the closest homologs of AtPCOs.

### Promoter analysis

One kb of genomic sequences upstream of each PCO gene translation start position (ATG) was obtained either through the phytozome https://phytozome.jgi.doe.gov/pz/portal.html) or ensemble (https://www.ensembl.org/index.html) portals. When annotated, the 5’ UTR region was also included in the analysis. The presence of HRPE (Gasch et al., 2016) cis-regulatory sequences was assessed using the FIMO package of MEME-suit 5.1.1 (Bailey et al., 2009). For each promoter, the number and position of HRPE elements was retrieved and noted in **Supplemental Table S3** and **Supplemental File S2**, respectively.

### Phylogenetic analysis

Phylogenetic analysis was performed using MEGAX (Kumar *et al*., 2018), by applying the Maximul Likelihood method and a JTT matrix-based model (Jones et al. 1992). To generate the phylogenetic tree, NCO protein sequences from different species were aligned using the MUSCLE algorithm (Edgar, 2004) and the initial trees were obtained by applying Neighbor-Join and BioNJ algorithms. The tree with the highest log-likelihood was selected and the bootstrap analysis (500 repeats) returned the percentage of trees in which the associated taxa clustered together.

### Assessment of gene expression levels

Total RNA extraction, DNAse treatment, cDNA synthesis and qRT-PCR analysis was performed as described previously (Kosmacz *et al*., 2015).

### PCO expression, purification and oxygen consumption assay

The coding sequences of *PCO4* and *PCO5* were cloned into pDEST17 vector (Thermo-Fisher Scientific) to bear a construct coding for PCOs tagged by a cleavable 6-His peptide at the N terminus. Protein expression and purification was performed as described previously (Weits *et al*., 2014). Oxygen consumption by PCO4 and PCO5 proteins was determined using an optical sensor (Presens, Germany) as a measure for enzyme activity as described before (Weits *et al*., 2014).

### Plant transformation

Stable transgenic plants expressing PCO4prom:GUS, 35S:GFP:PCO4 and 35S:GFP:PCO5GFP were obtained using the floral dip method (Clough & Bent, 1998). T0 seeds were screened for kanamycin resistance to identify independent transgenic plants. T3 generation plants were used for the experiments.

### Confocal imaging

For PCO-GFP imaging, the abaxial side of leaves from independently transformed plants (two weeks old) were analysed with a Leica DM6000B/SP8 confocal microscope (Leica Microsystems) using 488-nm laser light (20% laser transmissivity), PMT detection, and emission light was collected between 491 and 551 nm. Images were analysed and exported using the LAS X life science software (www.leica-microsystems.com), with an unchanged lookup table settings for each channel.

### Luciferase transactivation assay

Transactivation assays were performed using a dual luciferase assay based on *Renilla reniformis* and *Photinus pyralis* luciferase enzymes. A 31 nt long aequence containing the HRPE element was retrieved from the LBD41 promoter (−364 to −331 from the initial ATG), repeated five times in tandem and fused to a minimal 35S promoter (Supplementary Information File S1). This sequences was synthesized by Geneart (Thermo Fisher Scientific), inserted into pENTR/D-topo (Thermo Fisher Scientific) and recombined into the *pGREEN800LUC* plasmid (Hellens *et al*., 2005) using LR clonase mix II (Thermo Fisher Scientific) to generate a reporter vector 5xHRPE:PpLuc. To evaluate the effect of PCO proteins on RAP2.12-mediated activation of the 5xHRPE promoter, the effector plasmids were produced by recombining the CDS of PCOs, GFP and RAP2.12 from pENTR-D/TOPO into p2GW7 (Karimi *et al*., 2002). Mesophyll protoplasts were prepared and transformed following the protocol by (Yoo et al. 2007) using three micrograms of each plasmid. Proteins were extracted from protoplasts after a 16h incubation in WI medium using 100 μl of 1 × passive lysis buffer (Promega). Luciferase activities were measured using the Dual Luciferase Reporter Assay kit (Promega) with a Glomax 20/20 (Promega).

### Quantification of ethanol production

One-week old Arabidopsis seedlings were peeled from vertical plates and incubated for 12 h in 1 ml of liquid half-strength MS medium (pH 5.8) supplemented with 2% sucrose (w/v) with shaking. At the end of the light phase of the day, seedlings were treated with normoxia or anoxia (>0.01% O_2_ V/V) for 12 h. At the end of the treatment, the medium was collected and the ethanol release per milligram of fresh weight of plant material was measured as described by Licausi *et al*. (2010).

## Supplemental data

Figure S1. Developmental map of *AtPCO3* expression.

Figure S2. Developmental map of *AtPCO4* expression.

Figure S3. Developmental map of *AtPCO5* expression.

Figure S4. Occurrence of NCO-like sequences in Chytrid species.

Figure S5. Genotyping of *pco4, pco5* and *pco4pco5* mutants.

Table S1. Relative mRNA levels of PCO genes in Arabidopsis seedlings subjected to hypoxia.

Table S2. List of PCO sequences encoded in the genome of angiosperm species for which the hypoxic transcriptome is available.

Table S3. List of PCO-like sequences identified in the genome or transcriptome of species representing evolutionary steps towards the establishment of the angiosperm taxon.

Table S4. List of NCO-like sequences identified in the chytrid group of the fungal kingdom.

Table S5. Relative mRNA levels of PCO genes in different species subjected to hypoxia.

Table S6. Identification of nuclear localization sequences within the Arabidopsis PCO proteins

Table S7. Relative mRNA levels of anaerobiosis core-response genes in *pco* Arabidopsis.

Table S8. List of primers used in this study.

Table S9. List of DNA vectors used in this study.

File S1. Occurrence of HRPE element within 5’ upstream genomic sequences of PCO genes

File S2. Multialignment used to generate the phylogenetic tree used in Fig. 1C.

## Literature cited

Altschul SF, Gish W, Miller W, Myers EW, Lipman DJ (1990) Basic local alignment search tool. J. Mol. Biol. 215: 403–410.

Bachmair A, Finley D, Varshavsky A (1986) In vivo half-life of a protein is a function of its amino-terminal residue. Science 234: 179–186.

Bailey IL, Boden M, Buske FA, Martin Frith M, CE, Clementi L, Ren J, Li WW, Noble WS (2009) MEME SUITE: Tools for Motif Discovery and Searching – PubMed. Nucleic Acids Res 37: 202–208

Bailey-Serres J, Colmer TD (2014) Plant tolerance of flooding stress – recent advances. Plant, Cell & Environment 37: 2211–2215.

Chang Y, Wang S, Sekimoto S, Aerts AL, Choi C, Clum A, LaButti KM, Lindquist EA, Ngan CY, Ohm RA, et al (2015) Phylogenomic Analyses Indicate that Early Fungi Evolved Digesting Cell Walls of Algal Ancestors of Land Plants. Genome Biol Evol 7: 1590–1601

Chung HS, Wang SB, Venkatraman V, Murray CI, Van Eyk JE (2013) Cysteine oxidative post-translational modifications: emerging regulation in the cardiovascular system. Circulation research 112: 382–392.

Clough SJ, Bent AF (1998) Floral dip: a simplified method for Agrobacterium-mediated transformation ofArabidopsis thaliana. The Plant Journal 16: 735–743.

Dalle Carbonare LD, White MD, Shukla V, Francini A, Perata P, Flashman E, Sebastiani L, Licausi F (2019) Zinc Excess Induces a Hypoxia-Like Response by Inhibiting Cysteine Oxidases in Poplar Roots. Plant physiology, 180(3), 1614–1628.

Edgar RC (2004) MUSCLE: multiple sequence alignment with high accuracy and high throughput. Nucleic Acids Research 32: 1792–1797.

Gasch P, Fundinger M, Müller JT, Lee T, Bailey-Serres J, Mustroph A (2016) Redundant ERF-VII Transcription Factors Bind to an Evolutionarily Conserved cis-Motif to Regulate Hypoxia-Responsive Gene Expression in Arabidopsis. The Plant Cell 28: 160–180.

Gibbs DJ, Lee SC, Isa NM, Gramuglia S, Fukao T, Bassel GW, Correia CS, Corbineau F, Theodoulou FL, Bailey-Serres J et al (2011) Homeostatic response to hypoxia is regulated by the N-end rule pathway in plants. Nature 479: 415–418.

Gibbs DJ, Isa NM, Movahedi M, Lozano-Juste J, Mendiondo GM, Berckhan S, Marín-de la Rosa N, Vicente Conde J, Sousa Correia C, Pearce SP et al (2014) Nitric Oxide Sensing in Plants Is Mediated by Proteolytic Control of Group VII ERF Transcription Factors. Molecular Cell 53: 369–379.

Gibbs DJ, Tedds HM, Labandera AM, Bailey M, White MD, Hartman S, Sprigg C, Mogg SL, Osborne R, Dambire C, Boeckx T, Paling Z, Voesenek LACJ, Flashman E, Holdsworth MJ (2018) Oxygen-dependent proteolysis regulates the stability of angiosperm polycomb repressive complex 2 subunit VERNALIZATION 2. Nat Commun. 9: 5438.

Giuntoli B, Lee SC, Licausi F, Kosmacz M, Oosumi T, van Dongen JT, Bailey-Serres J, Perata P (2014) A Trihelix DNA Binding Protein Counterbalances Hypoxia-Responsive Transcriptional Activation in Arabidopsis. PLoS Biol 12: e1001950.

Giuntoli B, Shukla V, Maggiorelli F, Giorgi FM, Lombardi L, Perata P, Licausi F (2017) Age-dependent regulation of ERF-VII transcription factor activity in Arabidopsis thaliana. Plant Cell Environ. 40: 2333–2346

Graciet E, Mesiti F, Wellmer F (2010) Structure and evolutionary conservation of the plant N-end rule pathway. The Plant Journal 61: 741–751.

Graciet E, Wellmer F (2009) The plant N-end rule pathway: structure and functions. Trends in Plant Science 15: 447–453.

Hellens RP, Allan AC, Friel EN, Bolitho K, Grafton K, Templeton MD, Karunairetnam S, Gleave AP, Laing WA (2005) Transient expression vectors for functional genomics, quantification of promoter activity and RNA silencing in plants. Plant Methods 1: 13–13.

Henze A, Acker T (2010) Feedback Regulators of Hypoxia-Inducible Factors and Their Role in Cancer Biology. Cell cycle 14: 2749–2763

Holdsworth MJ, Gibbs DJ (2020) Comparative Biology of Oxygen Sensing in Plants and Animals. Curr Biol 30: R362–R369

Hu RG, Sheng J, Qi X, Xu Z, Takahashi TT, Varshavsky A (2005) The N-end rule pathway as a nitric oxide sensor controlling the levels of multiple regulators. Nature 437: 981–986.

Jones DT, Taylor WR, Thornton JM (1992) The Rapid Generation of Mutation Data Matrices From Protein Sequences – PubMed. Comput Appl Biosci 8: 275–282

Kagami M, Miki T, Takimoto G (2014) Mycoloop: Chytrids in aquatic food webs. Front Microbiol 5: 166

Karimi M, Inzé D, Depicker A (2002) GATEWAY^®^ vectors for Agrobacterium-mediated plant transformation. Trends in Plant Science 7: 193–195.

Kerpen L, Niccolini L, Licausi F, van Dongen JT, Weits DA (2019) Hypoxic Conditions in Crown Galls Induce Plant Anaerobic Responses That Support Tumor Proliferation. Front Plant Sci. 10:56.

Kosmacz M, Parlanti S, Schwarzlaender M, Kragler F, Licausi F, Van Dongen JT (2015) The stability and nuclear localization of the transcription factor RAP2.12 are dynamically regulated by oxygen concentration. Plant, Cell & Environment 38: 1094–1103.

Kumar S, Stecher G, Li M, Knyaz C, Tamura K (2018) MEGA X: Molecular Evolutionary Genetics Analysis across Computing Platforms | Molecular Biology and Evolution. Mol Biol Evol 35:1547–1549

Labandera AM, Tedds HM, Bailey M, Sprigg C, Etherington RD, Akintewe O, Kalleechurn G, Holdsworth MJ, Gibbs DJ (2020) The PRT6 N-degron pathway restricts VERNALIZATION 2 to endogenous hypoxic niches to modulate plant development. New Phytol. doi: 10.1111/nph.16477.

Licausi F, van Dongen JT, Giuntoli B, Novi G, Santaniello A, Geigenberger P, Perata P (2010) HRE1 and HRE2, two hypoxia-inducible ethylene response factors, affect anaerobic responses in Arabidopsis thaliana. Plant J 62: 302–15

Licausi F, Kosmacz M, Weits DA, Giuntoli B, Giorgi FM, Voesenek LACJ, Perata P, van Dongen JT (2011) Oxygen sensing in plants is mediated by an N-end rule pathway for protein destabilization. Nature 479: 419–422.

Liu YY, Yang KZ, Wei XX, Wang XQ (2016) Revisiting the phosphatidylethanolamine-binding protein (PEBP) gene family reveals cryptic FLOWERING LOCUS T gene homologs in gymnosperms and sheds new light on functional evolution. New Phytol 212: 730–744

Masson N, Keeley TP, Giuntoli B, White MD, Puerta ML, Perata P, Hopkinson RJ, Flashman E, Licausi F, Ratcliffe PJ (2019) Conserved N-terminal cysteine dioxygenases transduce responses to hypoxia in animals and plants. Science 365: 65–69

McCoy JG, Bailey LJ, Bitto E, Bingman CA, Aceti DJ, Fox BG, Phillips GN (2006) Structure and mechanism of mouse cysteine dioxygenase. Proc Natl Acad Sci U S A 103: 3084–3089

Mustroph, A, Zanetti ME, Jang CJH, Holtan HE, Repetti PP, Galbraith DW, Girke T, Bailey-Serres J (2009) Profiling translatomes of discrete cell populations resolves altered cellular priorities during hypoxia in Arabidopsis. Proceedings of the National Academy of Sciences 106: 18843–18848.

Paul MV, Iyer S, Amerhauser C, Lehmann M, van Dongen JT, Geigenberger P (2016) RAP2.12 oxygen sensing regulates plant metabolism and performance under both normoxia and hypoxia. Plant Physiology. 172:141–53.

Puerta ML, Shukla V, Dalle Carbonare L, Weits DA, Perata P, Licausi F, Giuntoli B (2019) A Ratiometric Sensor Based on Plant N-Terminal Degrons Able to Report Oxygen Dynamics in Saccharomyces cerevisiae. JMB 431: 2810–2820

Reddie KG, Carroll KS (2008) Expanding the functional diversity of proteins through cysteine oxidation. Current Opinion in Chemical Biology 12: 746–754.

Reski R, Abel WO (1985) Induction of budding on chloronemata and caulonemata of the moss, Physcomitrella patens, using isopentenyladenine. Planta 165: 354–358.

Riber W, Müller JT, Visser EJ, Sasidharan R, Voesenek LA, Mustroph A (2015) The Greening after Extended Darkness1 Is an N-End Rule Pathway Mutant with High Tolerance to Submergence and Starvation. Plant Physiology 167: 1616–1629.

Santaniello A, Loreti E, Gonzali S, Novi G, Perata P (2014) A reassessment of the role of sucrose synthase in the hypoxic sucrose-ethanol transition in Arabidopsis. Plant, Cell & Environment 37: 2294–2302.

Shukla V, Lombardi L, Iacopino S, Pencik A, Novak O, Perata P, Giuntoli B, Licausi F (2019) Endogenous Hypoxia in Lateral Root Primordia Controls Root Architecture by Antagonizing Auxin Signaling in Arabidopsis. Mol Plant 12: 538–551

Tasaki T, Sriram SM, Park KS, Kwon YT (2012) The N-End Rule Pathway. Annual review of biochemistry 81: 261–289.

Taylor TN, Remy W, Hass H (1994) Allomyces in the Devonian. Nature 367: 601

Weits DA, Giuntoli B, Kosmacz M, Parlanti S, Hubberten HM, Riegler H, Hoefgen R, Perata P, van Dongen JT and Licausi F (2014) Plant cysteine oxidases control the oxygen-dependent branch of the N-end-rule pathway. Nat Commun 5.

Van Dongen JT and Licausi F (2015) Oxygen Sensing and Signaling. Annu Rev Plant Biol 66: 345–67

Weits DA, Kunkowska AB, Kamps NCW, Portz KMS, Packbier NK, Nemec Venza Z, Gaillochet C, Lohmann JU, Pedersen O, van Dongen JT, Licausi F (2019) An apical hypoxic niche sets the pace of shoot meristem activity. Nature 569: 714–717.

Weits DA, van Dongen JT and Licausi F (2020) Molecular oxygen as a signaling component in plant development. New Phytol. doi:10.1111/nph.16424.

White MD, Klecker M, Hopkinson RJ, Weits DA, Mueller C, Naumann C, O’Neill R, Wickens J, Yang J, Brooks-Bartlett JC, et al. (2017) Plant cysteine oxidases are dioxygenases that directly enable arginyl transferase-catalysed arginylation of N-end rule targets. Nat Commun 8: 14690.

White MD, Kamps JJAG, East S, Taylor KLJ, Flashman E (2018) The Plant Cysteine Oxidases from Arabidopsis thaliana are kinetically tailored to act as oxygen sensors. JBC 293, 11786–11795.

Yoo SD, Cho YH, Sheen J (2007) Arabidopsis mesophyll protoplasts: a versatile cell system for transient gene expression analysis. Nat. Protocols 2: 1565–1572.

